# Transgenesis in the acoel worm *Hofstenia miamia*

**DOI:** 10.1101/2021.07.21.453044

**Authors:** Lorenzo Ricci, Mansi Srivastava

## Abstract

The acoel worm *Hofstenia miamia*, which can replace tissue lost to injury via differentiation of a population of stem cells, has emerged as a new research organism for studying regeneration. To enhance the depth of mechanistic studies in this system, we devised a protocol for microinjection into embryonic cells that resulted in stable transgene integration into the genome and generated animals with tissue-specific fluorescent transgene expression in epidermis, gut, and muscle. We demonstrate that transgenic *Hofstenia* are amenable to the isolation of specific cell types, detailed investigations of regeneration, tracking of photoconverted molecules, and live imaging. Further, our stable transgenic lines revealed new insights into the biology of *Hofstenia*, unprecedented details of cell morphology and the organization of muscle as a cellular scaffold for other tissues. Our work positions *Hofstenia* as a powerful system with unparalleled tools for mechanistic investigations of development, whole-body regeneration, and stem cell biology.

## Introduction

New research organisms hold the potential to reveal novel biological insights into phenomena that cannot be studied via established model systems. Numerous animal species have emerged as new systems for laboratory research, *e*.*g*., short-lived killifish for studying aging, hardy water bears for studying tolerance to extreme environments, or the anemone *Aiptasia* for studying coral bleaching (Lehnert, Burriesci and Pringle, 2012; Goldstein, 2018; Hu and Brunet, 2018). The lack of genetic tools in emerging systems can limit the depth of molecular and cellular insight, however, advances in genome-scale sequencing technology and tools for genetic manipulations (*e*.*g*., via CRISPR/CAS9) are releasing this constraint (Ikmi *et al*., 2014; Wudarski *et al*., 2017; Minor *et al*., 2019). Here, we developed a method for transgenesis in the acoel worm, *Hofstenia miamia* (Corrêa, 1960), which has emerged as a new research organism for studying regeneration and stem cell biology.

In contrast to the limited regeneration capacities of vertebrates, many invertebrate species regenerate robustly, reconfiguring entire body axes and replacing virtually any missing cell type. Notably, some of these highly regenerative invertebrates also harbor large populations of multipotent or pluripotent stem cells as adult animals (Hemmrich *et al*., 2012; Gahan *et al*., 2016; Wang, Wagner and Reddien, 2018; Kassmer, Langenbacher and De Tomaso, 2020). *Hofstenia* is capable of this “whole-body” regeneration and harbors adult stem cells referred to as “neoblasts”. A sequenced genome, transcriptomic resources, and robust and systemic RNA interference (RNAi) have enabled functional studies of regeneration and stem cells in *Hofstenia* (Srivastava *et al*., 2014; Raz *et al*., 2017; Gehrke *et al*., 2019; Ramirez, Loubet-Senear and Srivastava, 2020), and we sought to leverage the biology of this system further by developing tools for transgenesis.

*Hofstenia miamia, a*.*k*.*a*. the three-banded panther worm, belongs to an enigmatic phyletic lineage, the Xenacoelomorpha (comprising acoels, xenoturbellids and nemertodermatids), which is the likely sister-lineage to all other animals with bilateral symmetry (**Fig. 1A**), *i*.*e*., bilaterians (Cannon *et al*., 2016). Thus, studies in acoels are important for understanding the evolution of major features of bilaterians, such as anterior-posterior and dorsal-ventral axes, mesoderm, and a centralized nervous system. Moreover, this phylogenetic position places *Hofstenia* as evolutionarily distantly related to other regenerative model systems such as cnidarians, planarians, and sea squirts. Therefore, development of more genetic tools in this system has the potential to advance the study of an important yet understudied animal group as well as to inform the evolution of pathways for development, regeneration, and stem cell biology.

**Figure 1:**
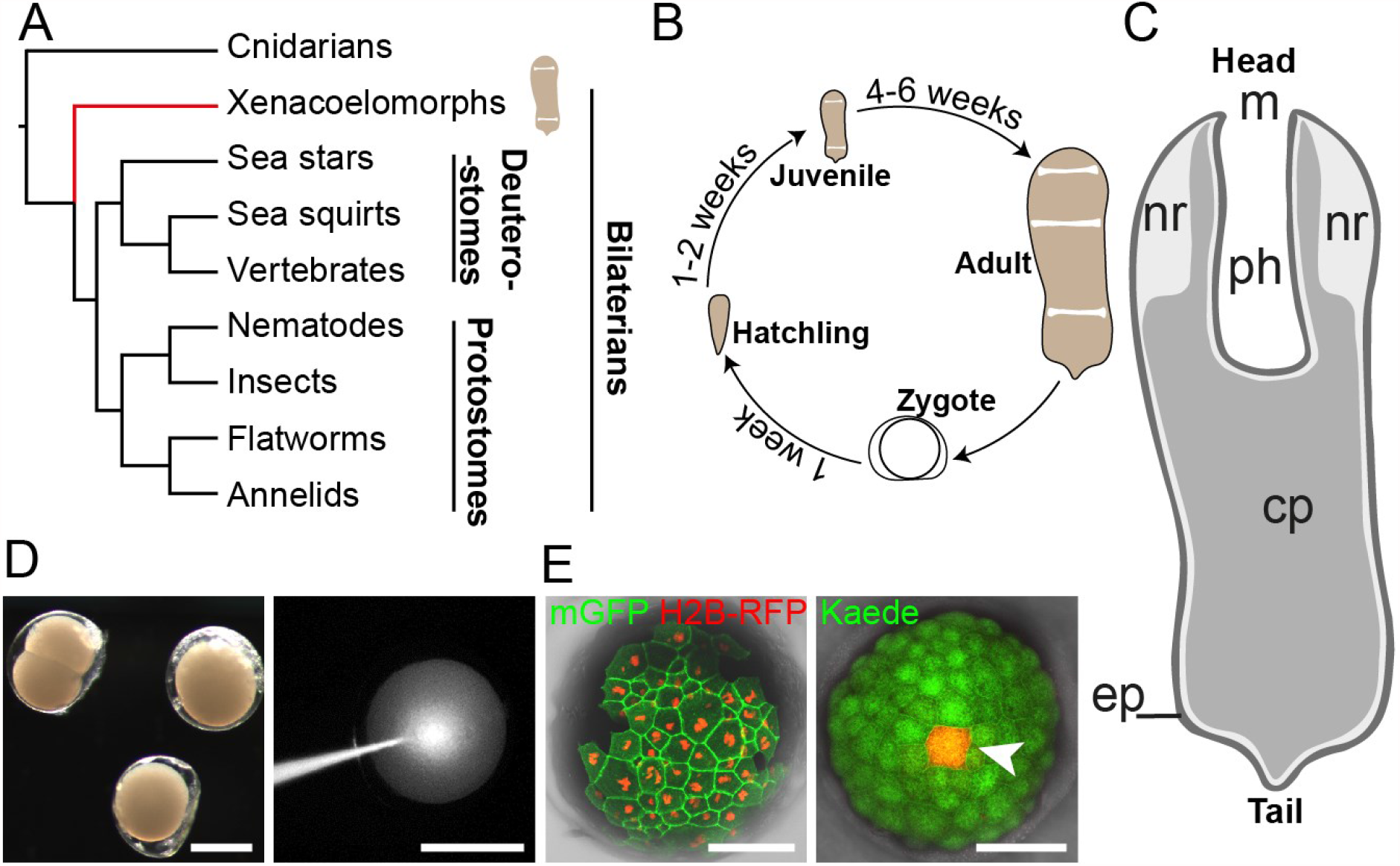
*Hofstenia miamia* is a highly regenerative species with accessible and manipulable embryos. **(A)** Simplified phylogenetic tree of metazoans showing placement of *Hofstenia* within xenacoelomorphs (red line). (**B**) An adult worm with schematic drawing showing major structures. (**C**) Life cycle of *Hofstenia miamia* in lab conditions. (**D**) *Hofstenia* embryos are amenable to microinjections. **Left**, live embryos at 2-cell and zygote stages; **right**, injection of FL-Dextran into a zygote. Scale bars, 200 µm. (**E**) Translation of foreign mRNAs. **Left**, detection of H2B-RFP and meGFP, 1.5 days after single blastomere injection of a 2-cell stage embryo; **right**, single cell photoconversion of Kaede (white arrowhead), 3 days after injection of Kaede mRNA into a zygote. Scale bars, 100 µm.

*Hofstenia* are hermaphroditic marine worms (Achatz *et al*., 2013; Jondelius, Raikova and Martinez, 2019) that produce accessible embryos, which develop and hatch into juveniles, which become sexually mature in 4-6 weeks (**Fig. 1B**). Given our ability to close the life cycle in the lab, we targeted the embryos of *Hofstenia* to develop tools for microinjection into early embryonic cells. First, we confirmed that *Hofstenia* embryonic cells were able to translate exogenous mRNAs coding for fluorescent proteins. Then, taking advantage of the recently sequenced genome of *Hofstenia*, we cloned the promoter regions of various genes, including markers of muscle (*troponin*), digestive cells (*saposin*), and anterior epidermis (a *Hofstenia*-specific gene we refer to as *epiA*), in a backbone designed for meganuclease-assisted transgenesis (Renfer *et al*., 2010). We generated multiple, stable transgenic lines, with strong, tissue-specific fluorescence. The resulting strains revealed, at high resolution, the morpho-anatomy of putatively ectodermal, mesodermal, and endodermal cells types in *Hofstenia*. Additionally, these animals were amenable to studies of regeneration, to efficient fluorescence activated cell sorting (FACS), and to photoconversion-based assays. Overall, our work demonstrates that *Hofstenia* is well-suited to *in vivo* studies and transgenesis, and is now accessible to many downstream applications for future studies of organogenesis, regeneration, and embryogenesis.

## Results

### Embryonic microinjection and transgenesis in Hofstenia

We first sought to deliver nucleic acids into embryonic cells in *Hofstenia*. The life cycle of *Hofstenia* is completed within two months, under ideal laboratory conditions (**Fig. 1B, see *animal culture* in methods**), and adults produce an average of four embryos a day that hatch within 7-8 days as juvenile worms, resembling the overall morphology of the adults (**Fig. 1C**). Embryos are deposited on the substrate as zygotes, with each embryo enveloped by a transparent egg “shell”. We found that this tough outer covering of the embryo can be penetrated by a sharp quartz microinjection needle (**Fig. 1D**). As a first step toward transgenesis, we assessed the ability of *Hofstenia* cells to receive and effectively translate foreign mRNA molecules. We injected single blastomeres, at early embryonic stages (2- and 4-cell embryos) with *in vitro* synthesized mRNAs encoding fluorescent proteins (FPs) with a range of green and red emission spectra. Among these, eGFP, Kaede, mRFP, and TagRFP-T offered optimal brightness and stability. We found that subcellular localization signals added to the injected mRNA (nuclear and plasma membrane) targeted the FPs to the appropriate cellular compartments. Additionally, we found that the green fluorescent protein Kaede could be efficiently converted, at single cell resolution, to emanate red fluorescence in *Hofstenia* embryos injected with Kaede-encoding mRNA (**Fig. 1E**).

### Establishment of stable transgenic lines

We next aimed to generate transgenic fluorescent reporter lines in order to label specific cell types in *Hofstenia*. We identified genes encoded in the *Hofstenia* genome that, 1) either based on homology or on known fluorescent *in situ* hybridization (FISH) data were likely to show cell type specific expression, and 2) were expressed at high levels in transcriptome sequencing data and in FISH experiments. This yielded three genes as candidates for generating transgenic *Hofstenia*: a *Hofstenia*-specific gene expressed in epidermal cells in the anterior (*epidermis anterior*; *epiA*), a homolog of *saposin* expressed in the gut (*saposin, sap*), and a homolog of *troponin* expressed in muscle (*troponin, tnn*) (**Supp. Fig. 1**). Given that epidermis, gut, and muscle are usually derived from ectodermal, endodermal, and mesodermal tissues, respectively, in bilaterians, these genes are putative markers of these three tissue types in *Hofstenia*. We cloned regulatory regions of the candidate genes into a backbone plasmid with restriction sites for the I-SceI endonuclease flanking the transgenic cassette (**Fig. 2A**), a system that has been used successfully in multiple systems including cnidarians, protostomes, and vertebrates (Thermes *et al*., 2002; Deschet, Nakatani and Smith, 2003; Ogino, McConnell and Grainger, 2006; Renfer *et al*., 2010; Minor *et al*., 2019).

**Figure 2:**
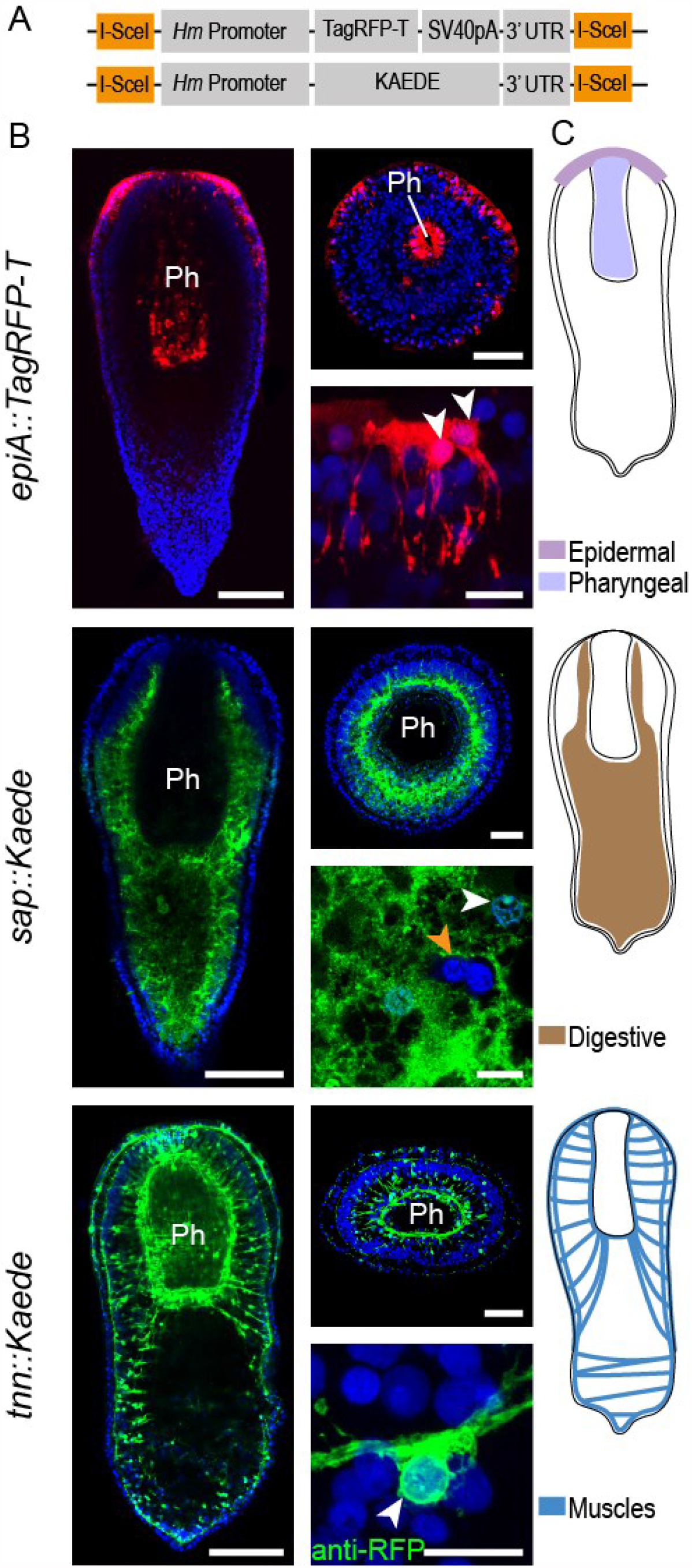
Transgene expression reveals structure and cellular anatomy of epidermal, digestive, and muscle tissues. (**A**) Schematic of the two transgenic cassettes used in this study. Purple box indicates a recognition site for the Meganuclease enzyme. (**B**) Expression of fluorescent reporter genes under control of tissue-specific *Hofstenia* promoters (name of the transgene is indicated on the left) in fixed, juvenile worms. **Left column**: whole worm, mid-body confocal section, as observed in G1 and G2 animals. **Right column, top**: cross section, at the pharynx level; **bottom**: high magnification images of transgene-expressing cells. DAPI (nuclei, blue); Kaede (green); TagRFP-T (red); except for *tnn::Kaede* right bottom picture, anti-RFP : immunodetection of TagRFP-T. White arrowheads point to nuclei of transgene-expressing cells; orange arrowhead indicate nuclei of cells embedded in digestive tissue and not expressing the transgene. Scale bars: whole animal, 100 µm; cross section, 50 µm; close up, 10 µm. (**C**) Schematics depicting the transgene expression as observed in G1 and G2 generations. *Ph, Pharynx*. Images and sketches of whole worms are all oriented anterior side up.

After determining the range of DNA concentrations offering high transgene expression and low lethality (**Table1**), we co-injected large batches of *Hofstenia* embryos at the zygote and 2-cell stages with the I-SceI enzyme and plasmid DNA at this optimal concentration (10-25 ng/μl). Next, we screened embryos regularly to detect fluorescence. We found that around 40% of the embryos exhibited fluorescence in at least one cell during development. Upon hatching, G0 worms injected with and expressing the *tnn* or the *epiA* constructs showed mosaic expression of the transgene. Notably, embryos injected with the *sap* construct hatched into worms with their entire digestive tissue labeled (**Supp. Fig. 1**), consistent with the presence of a syncytial digestive system, which has been reported for some acoels (Smith, 1981; Gavilán *et al*., 2019). Shortly after hatching, we sorted G0 animals based on the presence of transgene expression and the intensity of fluorescence: only those exhibiting strong fluorescence and the largest area of expression were raised to adulthood and screened regularly for continued transgene expression. Putative adult transgenic worms were then mated to wild type animals or to each other, when possible. Screening of fluorescence patterns in G1 animals suggests that at least 1% (and up to 3%) of the injected G0 animals experienced stable integration of the transgene into the genome with transmission to the germline (**Table1**). G2 embryos and animals tended to show higher levels of fluorescence, likely due to a higher number of transgene copies, and/or due to homozygous transgenic loci. For this study, we focused on three transgenic lines: *epiA::TagRFP-T, tnn::Kaede* and *sap::Kaede*. Additionally, we generated *tnn::TagRFP-T* and *sap::TagRFP-T* lines to facilitate imaging in fixed specimens using an anti-TagRFP antibody.

**Table 1:** Efficiency of transgenesis. The survival rate for DNA concentration ranges superior to 25 % was calculated from various batches of at least 50 injected embryos each. (*) Calculated from 2 different batches, including zygotes and 2-cell stage injected embryos. (n = 3/97 and n = 2/182).

| Injected DNA concentration (ng/ul) | Survival Rate (%) | G0 expression (%) | Germline Transmission (%) |
| --- | --- | --- | --- |
| 10-25 | 90* | 36-57* | 1-3 * |
| 40-50 | 50-60 | ND | ND |
| 75-100 | 10-20 | ND | ND |
| >100 | 5-10 | ND | ND |

### Correspondence of transgene fluorescence and mRNA expression patterns

Overall, transgene expression driven by the promoters of the marker genes recapitulated the corresponding mRNA expression detected via FISH. In the case of *tnn* and *sap* lines, the transgene expression patterns in G1 animals (and subsequent generations) matched FISH results perfectly (**Fig. 2, Supp. Fig. 1B**), with the entire muscle and digestive compartments being labelled, respectively, at all life stages. In contrast, the *epiA* expression pattern varied across individuals, both in transgenic worms and in FISH experiments. Although an overall anterior-posterior gradient was observed in juvenile worms in the epidermis in addition to expression in the pharynx, the number and the size of labelled patches of cells were highly variable between animals in both FISH and transgenic animals (**Fig. 2, Supp. Fig. 1B**). As animals grew and matured, *epiA* mRNA continued to be expressed in the epidermis, but became increasingly restricted in the anterior and was detectable, by FISH, in adult worms in a ring of cells located peripherally to the mouth and in two ventrolateral lines (**Supp. Fig. 1B**). Additionally, during growth, pharyngeal expression of *epiA* was progressively lost, as a new region of expression appeared ventrally, in cells forming the gonopore, in both transgenic animals and FISH experiments (**Supp. Fig 1B, Supp. Fig. 3B**). In adult transgenic worms, the *epiA* expression pattern was similar to that detected via FISH, except for the absence of the two ventrolateral lines (**Supp. Fig. 3B**).

### Organ structure and cellular anatomy revealed by transgenic Hofstenia

Animals with fluorescently-labeled epidermis, gut, and muscle in hand, we utilized high resolution imaging to understand the structure of these tissues. In addition to studying fixed transgenic worms of G1 and G2 generations, we also utilized G0 animals, where mosaic labeling enabled us to visualize single cells.

*EpiA+* cells in the *Hofstenia* epidermis showed unusual morphology. Classically, epidermis in invertebrates forms an epithelial tissue, with a flat basal surface. *Hofstenia* cells instead exhibited a large number of thin processes, similar to those described in the other acoel species *Convoluta pulchra* (Tyler and Rieger, 1999), (approximately 10 µm in length and 0.5 µm in thickness), protruding at the basal end of the tissue and projecting internally (**Fig. 2, Supp. Fig. 2B, Supp. Video 1**). The apical surface of these epidermal cells showed abundant ciliature and appeared polygonal, but lacked a consistent stereotypical shape (Todt and Tyler, 2007) (**Supp. Fig. 2A, 2C**). In addition, *epiA* expression was also detected in subsets of cells forming the inner wall of the pharynx (**Fig. 2B, Supp. Fig. 2D, 5A**). These pharyngeal *epiA*+ cells appeared to be contiguous with the mouth epidermis and resembled the morphology of the epidermal *epiA*+ cells in the body wall (**Fig. 5B, 5E, Supp. Fig. 5A**).

*Sap*::*Kaede* animals displayed an extensive area of transgene expression along the anterior-posterior axis, from the head (surrounding the statocyst) down to the tip of the tail, in the interior of the worms (**Fig. 2B, Supp. Fig. 2E-H**). We also observed that *Sap-*cells were present within the mass of *sap+* tissue (**Fig. 2B, Supp. Fig. 2**), and, additionally, that large portions of *sap+* tissue were going through the pharyngeal sphincter to the pharynx lumen in fixed specimens, suggesting that *Hofstenia* is able to partially evaginate its digestive tissue (**Supp. Fig. 2G, 2H**).

*Tnn* transgenic lines revealed both global muscular architecture and the anatomy of individual muscle fibers at single cell resolution in *Hofstenia*. Individual muscle cells, regardless of their directionality, displayed overall similar morphology: an elongated main fiber displaying terminal ramifications, with a small cell body located in a slightly shifted position relatively to the axis of the fiber, consistent with observations based on FISH (Raz *et al*., 2017) (**Fig. 2B, Supp. Fig. 4A**).

We elaborate further on the properties of epidermal, digestive, and muscle tissue, their interactions, and the utility of the corresponding transgenic lines in the following sections.

### A toolkit enabled by transgenic animals

Following these first steps, we evaluated the amenability of our various transgenic lines to new approaches for future research. First, we aimed to sort and isolate live cells from G2 *tnn* transgenic animals, via fluorescence activated cell sorting (FACS). Using a single adult *tnn::Kaede* worm, we developed a protocol to identify and isolate Kaede+ cells (**Fig. 3A**). We found that the green fluorescence observed was specific to Kaede-containing animals as there was very low background fluorescence in wild type worms. We tested the specificity of the cell isolation procedure by applying a custom anti-Tropomyosin (TPM) antibody (Hulett, Potter and Srivastava, 2020) to our cell suspension. We observed that the Kaede-gated cells exhibited both green fluorescence and reactivity to anti-TPM, indicating successful and specific isolation of muscle cells (**Fig. 3B, supp Fig. 3A**). Satisfyingly, only a small fraction of the control Kaede-gated cells showed reactivity to the antibody, and none of them exhibited green fluorescence.

**Figure 3:**
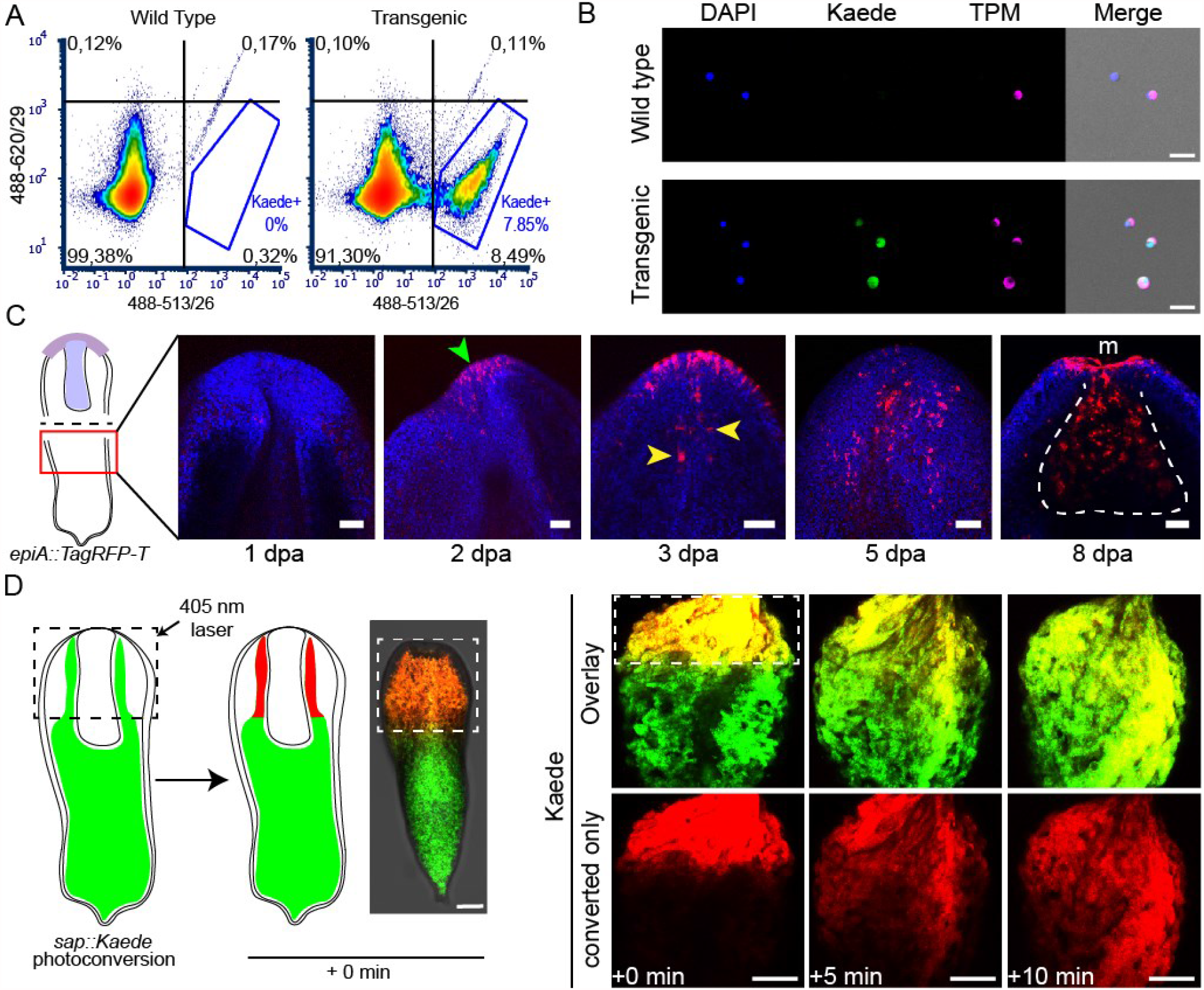
Transgenesis in *Hofstenia* enables a vast array of experimental tools for studying regeneration and stem cell biology. (**A, B**) Isolation of specific live cell populations using FACS on tnn::Kaede transgenic animals; (**A**) Fluorescence plots showing cell density for Kaede fluorescence (quadrant and Kaede gate percentages are calculated over a total of 76880 and 87157 singlets for wild type and transgenic, respectively); (**B**) anti-TPM ICC on FACS-isolated cells originating from the whole cell (wild type), and the Kaede+ (transgenic) populations. Scale bars, 20 µm. (**C**) Regeneration of anterior structures following worm bisection (see sketch on the left) in epiA::TagRFP-T transgenics. Regeneration time points are indicated under each image, as days post amputation (dpa). Green and yellow arrowheads point at de novo expression of TagRFP-T in epidermal and pharyngeal cells, respectively. White dotted line delineates the regenerated pharynx. M, mouth. Scale bars, 50 µm. (**D**) Photoconversion of Kaede-expressing cells in live sap::Kaede animals (dotted squares outline the photoconverted area); **left**, schematic of the experiment; **right**, quick expansion of the photoconverted Kaede area (red) over time in a live worm. Time elapsed since photoconversion is indicated at the bottom. Scale bars, 100 µm.

Next, to assess the value of transgenic *Hofstenia* for studying regeneration, we asked if transgenes would be re-expressed in new cells that form during regeneration. We took advantage of the anteriorly-restricted expression pattern of the *epiA* transgene and performed mid-body bisection of G1 animals to remove all anterior tissues including brain and pharynx. During regeneration, the anterior fragments, which lacked posterior/tail tissues, retained the *epiA* expression pattern as prior to the bisection (**Supp. Fig. 3B**). Posterior fragments, which lacked anterior/head tissues regained transgene expression over time, with expression first observable in the epidermis two days post amputation (**Fig. 3C, Supp. Fig. 3B**). Although we failed to detect *epiA+* pharyngeal cells immediately upon amputation in these fragments, expression of the transgene became visible in the newly forming pharynx three days post amputation.

Finally, we sought to determine if Kaede-expressing animals were amenable to photoconversion. Following embedding of *sap::Kaede* juveniles in agarose gel, we exposed them to 405 nm wavelength laser fluorescence, in order to partially photoconvert the digestive tissue. Within ten minutes following the photoconversion, we observed a rapid expansion of the region with red labeling, accompanied by a decrease in brightness for the same wavelength (**Fig. 3D**). This result, *i*.*e*. the spread of converted red fluorescent protein outside of the area that was targeted for conversion, confirms that the *Hofstenia* gut functions as a syncytium. Furthermore, together with our ability to photoconvert Kaede in single cells, this ability to study live adult worms suggests that photoconversion approaches can be utilized for cell tracking in *Hofstenia*.

### The structure and formation of musculature

The robust fluorescent labeling of muscle in the *tnn*::*Kaede* line provides an opportunity to study the formation of muscle cells during development and to determine the anatomy of musculature in detail. First, we fixed *tnn::Kaede* hatchlings in order to acquire fine details of muscle morphology in *Hofstenia*. Overall, we found that musculature is organized in four major interconnected compartments (**Fig. 4A**), consistent with previous observations (Todt, 2009). Starting from the outside of the body, we observed : *i)* an external layer of strictly longitudinal fibers with ramifications at their tips (the peripheral muscles) (**Fig. 4A, Supp. Fig. 4A**), running from the mouth towards the posterior, but not reaching the tip of the tail (**Fig. 4A, Supp. Fig. 4B**); *ii)* a tight network comprising longitudinal, circular and oblique fibers (the body wall muscles), also with ramifications, located beneath the epidermis, all along the A/P axis; *iii)* a set of highly ramified fibers projecting through the parenchyma (the parenchymal muscles) and connecting body wall muscles to each other or connecting body wall muscle to pharynx muscle (**Fig. 4A, Supp. Fig. 4C**) and *iv)* a dense, three-layered, basket-shaped network (**Fig. 4A, Supp. Fig. 4D, E**), of longitudinal and circular fibers delimiting the pharynx (the pharyngeal muscles).

**Figure 4:**
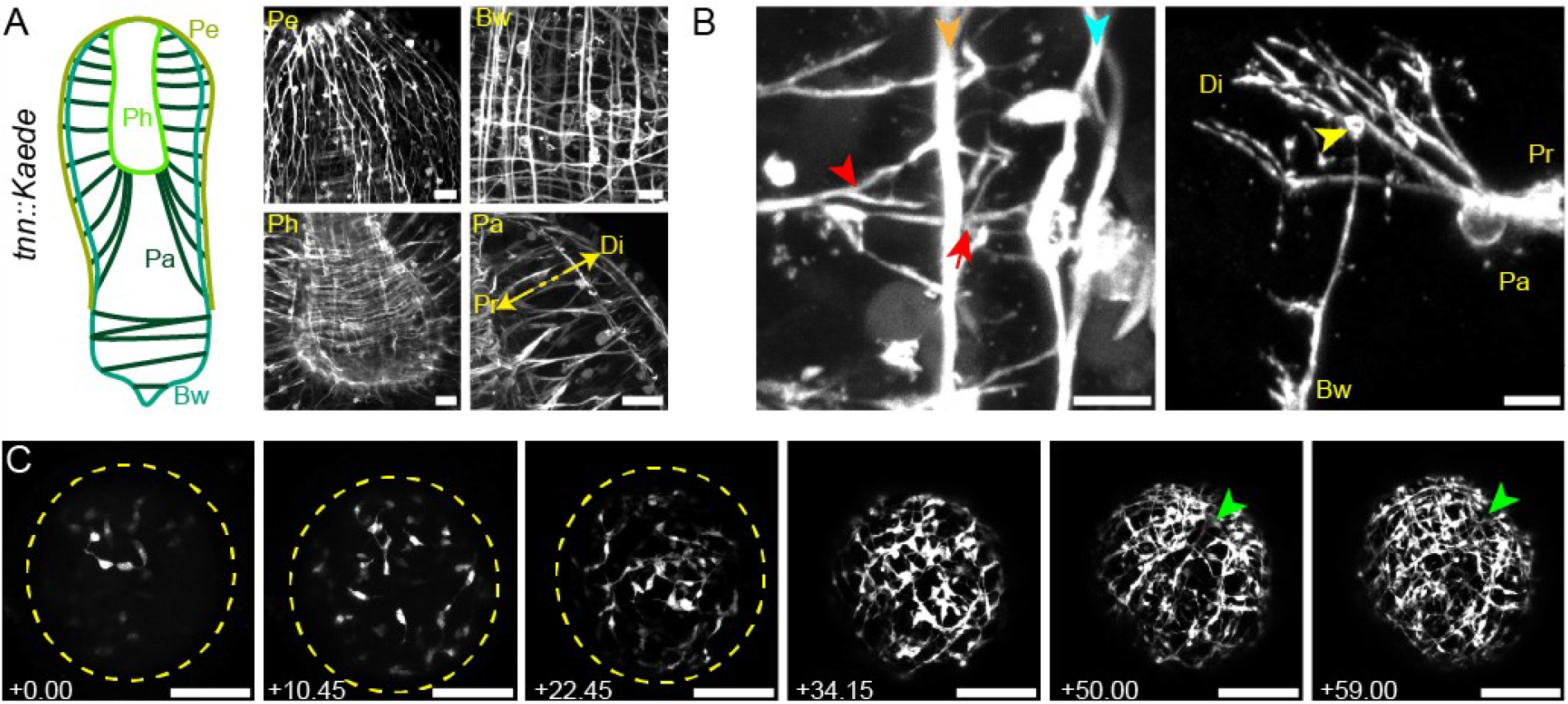
A muscle-specific reporter line reveals a robust network of interconnected muscle systems. (**A**) Major muscule structures of *Hostenia*. **Left**, sketch showing the overall organization of *Hofstenia* muscles. **Right**, close-up images of the different muscle categories. *Pe, peripheral; Bw, body wall; Pa, parenchymal; and Ph, pharyngeal muscles*. The double, yellow arrow indicates the proximal (*Pr*) and distal (*Di*) sides of the parenchymal muscles. Scale bars, 20 µm. (**B**) Muscle-muscle interactions. **Left**, proximal (red arrowhead) and distal (red arrow) ramifications of a parenchymal muscle fiber, connecting it to body wall (orange arrowhead) and peripheral (blue arrowhead) muscle fibers, respectively; **right**, overview of two orthogonal fibers connecting through ramified projections as observed in a mosaic, G0 animal. The yellow arrowhead indicates the thickened extremity of the body wall muscle fiber. *Pa, bw, single parenchymal and body wall muscle fibers, respectively, Pr, Di, proximal and distal sides of the parenchymal fiber, respectively*. Scale bars, left, 5 µm; right, 10 µm. (**C**) Birth and organization of the muscular network during a time-lapse movie of embryonic development. Left to right, Z-stack projections of the same embryo at 6 time points, ranging from 4 to 6 days after first embryonic cleavage. Yellow dashed lines outline the embryo at early stages of *tnn* expression; green arrowheads indicate the position of the future mouth. Scale bars, 100 µm.

Closer examination of the parenchymal muscle revealed a striking, interlocking feature of muscle in *Hofstenia*. In the anterior region of the animal, where the pharynx is located, parenchymal muscles contacted pharyngeal muscle proximally (towards the interior of the animal), and body wall and peripheral muscle distally (towards the periphery of the animal) (**Fig. 4B**). The contact between parenchymal muscle and body wall/peripheral muscle was mediated by multiple ramifications of the muscle fiber. As parenchymal muscle fibers approached body wall muscle, they formed ramifications, many of which made connections with the longitudinal body wall muscle. Some of these ramifications became further branched as they approach and made contact with the peripheral muscle. In the posterior, parenchymal muscles connected body wall muscle on opposite sides of the worm via ramifications similar to those observed in the anterior (**Supp. Fig. 4C**). Using 3D reconstruction tools, we were able to visualize more clearly how the two kind of fibers, parenchymal, and body wall (longitudinal) interact at their extremities through ramified interdigitations. In particular, 3D rendering of G1/G2 *tnn::Kaede* animals allowed us to clearly visualize the spatial arrangement of the parenchymal, body wall and peripheral muscles (**Supp. Videos 2A, 2B**). Additionally, G0 animals showed that the ramifications of a single parenchymal fiber interdigitate with those of a body wall fiber (**Supp. Videos 3A, 3B**), suggesting that parenchymal fibers do not randomly establish contact, but rather preferentially connect to ramifications at the tips of body wall muscle fibers.

Next, we developed approaches to live-image transgenic embryos. We focused on a subset of G2 transgenic worms that showed enhanced fluorescence relative to G0 and G1 animals. Such embryos started expressing Kaede around 3.5 days post laying, allowing *in vivo* time lapse imaging of the birth of muscle cells and the development of the musculature (**Fig. 4C, Supp. Video 4**). Early Kaede expression was first seen in a few fast-moving cells with round cell bodies, from which emanated short and labile plasma membrane projections. These cells increased in number due to rapid cell division (**Supp. Video 4**). At around 4 days post laying, these cells became more stationary and their projections increased in length, establishing contact with projections from other muscle cells and beginning to resemble contractile muscle fibers. Between 4.5 to 6 days post laying, muscle fibers progressively organized into a regular grid that formed the body wall muscle. Eventually, at around 6 days post laying, with only the body wall muscle visible, embryos started rotating within their capsules, thus, making time-lapse impossible. The structures formed by the peripheral, parenchymal and pharyngeal muscles could only be observed in later stages of development, suggesting that the body wall muscle network develops first during embryogenesis.

### A muscle scaffold for epidermal and digestive cells

The unique morphology of *Hofstenia* epidermis, with cells that project cytoplasmic processes toward the interior of the worm, the presence of a syncytial gut, and the interlocking organization of its muscular system, raised the question of the spatial relationship between these tissues. Taking advantage of our transgenic lines and of our custom made anti-TPM antibody, we performed dual labeling of muscle with epidermal, pharyngeal, and digestive cells, either by crossing G1 transgenics together, or by immunolabeling of muscles in *epiA::TagRFP-T* and *sap::TagRFP-T* transgenic lines.

In *epiA::TagRFP-T* X *tnn::Kaede* double transgenics (**Supp. Fig. 5A**) we observed, first, that both epidermal cells and peripheral muscles had their cell bodies located in the same plane, slightly distal relative to the plane defined by the peripheral muscle fibers. Second, we found that the epidermal processes emerged starting at the plane containing peripheral muscle fibers and extended proximally till they reached the body wall muscle grid (**Fig. 5A, D**). In the anterior two thirds of the animal, where parenchymal muscles connect to both pharyngeal and body wall muscles, epidermal processes were always associated with the distal-most ramifications of parenchymal muscle fibers, in the narrow space located between body wall and peripheral muscle fibers (**Fig. 5D**). In the posterior region of the worms, which lacks peripheral muscles, epidermal cell processes were not in contact with parenchymal fiber ramifications, but rather contacted the body wall grid alone (**Supp. Video 5**).

**Figure 5:**
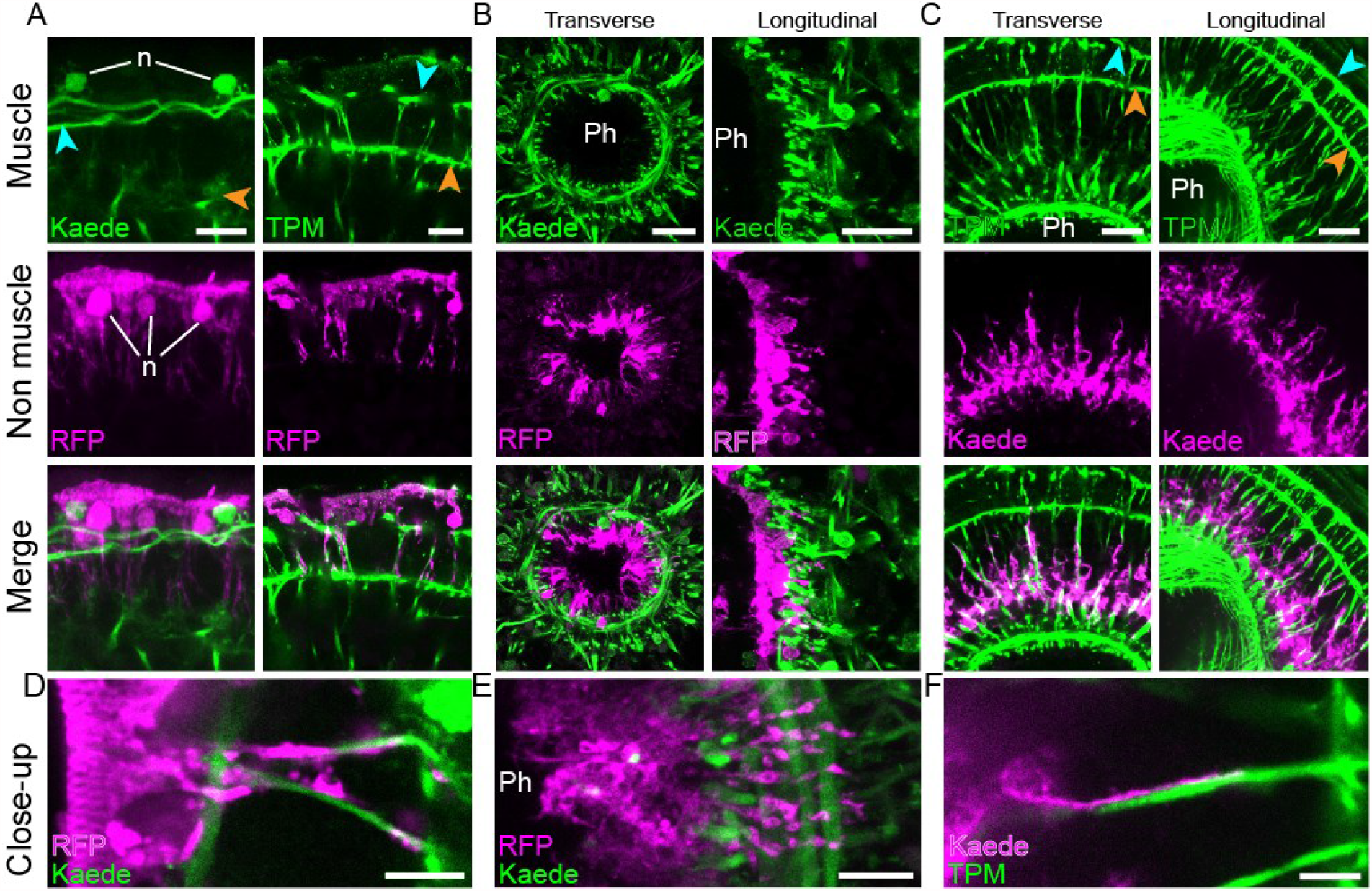
Muscle fibers closely contact epidermal, pharyngeal, and digestive cells. Dual labeling of muscles with epidermal (A, D), pharyngeal (B, E) and digestive (C, F) tissues. (**A**) Interaction of epidermal cells with peripheral muscle fibers (**left**) and ramifications of parenchymal muscle fibers (**right**). (**B**) Pharyngeal cells projecting ramifications through pharyngeal muscle net. Cross (**left**) and longitudinal (**right**) sections of the pharynx. (**C**) Digestive tissue projecting ramifications along parenchymal muscle fibers as seen in cross (**left**) and longitudinal (**right**) sections. (**D, E and F**) High magnification images showing the tight interaction between muscle fibers and epidermal, pharyngeal and digestive cell ramifications, respectively. Blue and orange arrowheads point at peripheral and body wall muscles, respectively. Images are in pseudo-colors; the fluorophore names are indicated in the left bottom corner (TPM, anti-Tropomyosin immunolabeling; TagRFP-T is abbreviated in RFP); except for TPM, fluorescent labeling originates from transgene expression. *Ph, pharynx lumen; n, nuclei*. Scale bars, A, 10 µm ; B and C, 20 µm ; D, E, F, 5 µm.

Given their shared morphological features with epidermal cells, we also looked at the interaction of pharyngeal *epiA+* cells with pharynx muscles (**Fig. 5B, E**). Although the high density of the pharyngeal muscle network did not allow clear observation of muscle fiber ramifications, we found that pharyngeal *epiA*+ cells exhibited narrow processes, similar to the epidermal cell processes, projecting through the net of pharyngeal muscles, towards the parenchymal space.

Because *Hofstenia* parenchymal muscles traverse a space that is occupied with the digestive syncytium, we interrogated the spatial arrangement between these two tissues. TPM immunolabeling in *sap::Kaede* animals revealed a close association of parenchymal muscle fibers with digestive tissue in both anterior and posterior regions of the animal (**Fig. 5C, F; Supp. Fig. 5B**). The syncytial gut exhibited on its distal edges thin, radial processes that aligned closely with parenchymal muscle fibers, reaching out to the body wall muscle grid. Together, these observations suggest that musculature, and parenchymal muscle in particular, has a prominent role in providing mechanical support, and potentially information to other tissues.

## Discussion

### Transgenesis in Hofstenia via random insertion

In this manuscript, we present the first successful attempt at establishing transgenesis in a xenacoelomorph species, the acoel *Hofstenia miamia*. Injection of a linearized transgenic cassette, comprising an FP-coding reporter gene flanked by promoter and 3’UTR regions of *Hofstenia* genes (**Fig. 2A**), resulted in the transient expression of the reporter gene in a large proportion of the injected embryos, and, subsequently in the stable integration of the transgene into the genome. With a rate of germline transmission of the transgene ranging between 1-3% of the injected animals (**Table 1**), we report that non-targeted transgenesis in *Hofstenia* has similar success rate to other invertebrate research organisms (Renfer *et al*., 2010; Backfisch *et al*., 2013; Wudarski *et al*., 2017). In particular, given the length of *Hofstenia* life cycle (**Fig. 1B**) we demonstrate the ability to generate stable G2 transgenic lines within 6 months.

The activity of the transgene promoters, recapitulated expression driven by their wild type counterparts, showing a high level of specificity of transgenesis in *Hofstenia*. The absence of *epiA::TagRFP-T* expression in the ventrolateral lines of adult worms (**Supp. Fig. 1B**,**3B**) represents the sole discrepancy detected between the transgenic and the wild type promoters. It is possible that the promoter we cloned did not include the regulatory elements needed for expression in this domain. We also demonstrate that *Hofstenia* transgenics are suitable for FACS-mediated isolation of specific live cell populations (**Fig. 3A**), prolonged live imaging of embryonic development (**Fig. 4C, Supp. Fig. 6**) and monitoring of the regeneration process (**Fig. 3B**). Furthermore, the use of Kaede as a reporter gene allowed us to differentially label a small portion of tissue and trace the photoconverted protein (**Fig. 3C**). Together, our results show the significant potential of transgenic *Hofstenia* for the study of acoel biology, and particularly for *in vivo* and live imaging applications.

**Figure 6:**
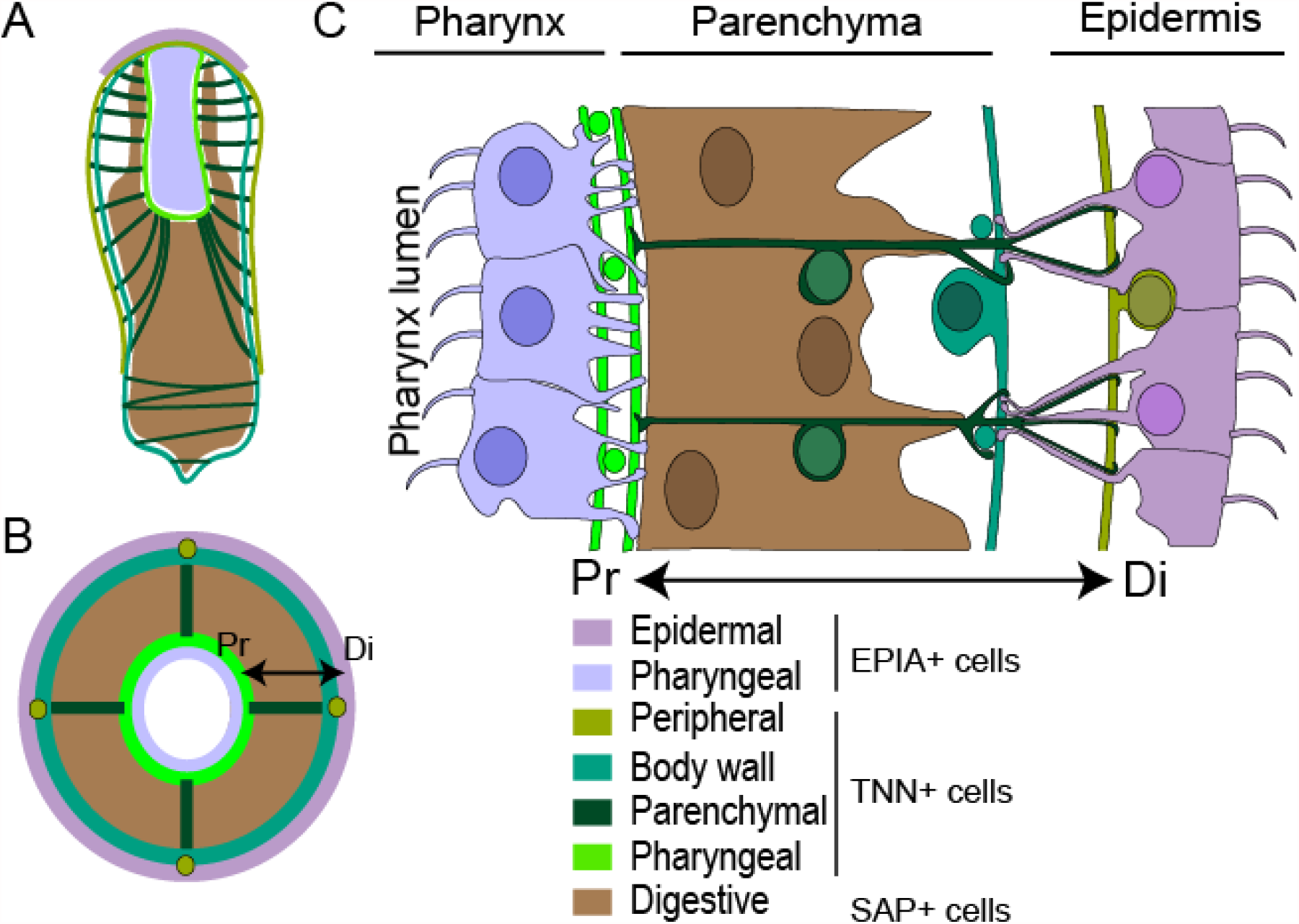
Anatomical model for *Hofstenia miamia*. Sketches of *Hofstenia* juvenile along a coronal (**A**) and a cross (**B**) section depicting the arrangement of epidermis, gut and muscles. (**C**) Drawing showing the detail of the cell type distribution as observed in a longitudinal section at the pharynx level. The double arrow points indicates the proximal (*Pr*) and distal (*Di*) ends of the parenchymal muscle.

### High resolution description of the musculature

The function of muscle cells in regeneration has recently been emphasized in planarians (Witchley *et al*., 2013; Scimone, Cote and Reddien, 2017; Tewari *et al*., 2019; Scimone *et al*., 2020), where they appear to guide progenitor cells to their correct location, notably by establishing a chemical coordinate system, through release of signaling molecules (Position Control Genes, PCGs; *e*.*g*., several *wnt, fz-4, bmp4, admp*).

Quite remarkably, despite the ancient divergence of acoels and planarians (550 mya), *Hofstenia* body wall muscles seem to share, at least partially, this instructive role as they express homologs of PCGs that are required for patterning of tissue during regeneration, and, as in planarians, PCG expression is triggered in muscle by regeneration (Raz *et al*., 2017; Ramirez, Loubet-Senear and Srivastava, 2020). Our results show a high degree of interconnection between the different muscle compartments (**Fig. 4**). In particular, parenchymal muscle appears central to global muscle organization, as it connects, pharyngeal muscle to body wall and peripheral muscle, making *Hofstenia* muscle an undisrupted, robust network. Therefore, we suggest that parenchymal muscle could participate in tissue renewal and/or regeneration 1) by facilitating the role of body wall muscles in those processes, *e*.*g*., by delivering positional and differentiation signals, or, 2) by providing physical support for stem cell homing and progenitor cell migration along the parenchymal fiber axis.

Transgenically-labeled epidermal cells revealed a unique morphology for this tissue in *Hofstenia*, with cytoplasmic processes projecting toward the interior of the animal. Parenchymal muscle fibers made close contacts with these epidermal processes, as well as with pharyngeal and digestive processes, suggesting that the muscular system provides mechanical support for these tissues (**Fig. 5-6**). In this regard, it would be interesting to investigate further the unique anatomy of *Hofstenia* and the role of its muscles, in particular by determining if and how muscle is associated with other tissues or cell types (*e*.*g*., stem cells, neurons, gland cells), and if, like planarians, acoel muscles are a source of extracellular matrix components (Cote, Simental and Reddien, 2019).

### Perspectives

For future mechanistic studies of regeneration and for understanding the diversity of regenerative and developmental mechanisms that emerged across evolution, it will be important to develop additional transgenic tools in *Hofstenia*. As an indication of the feasibility for more tool development in *Hofstenia*, we also report here CRISPR/CAS9 mediated genome editing in *Hofstenia* (**Supp. Fig. 10**). As a next step, we aim to develop targeted, and conditional transgenesis in order to minimize the biases of foreign DNA integration in the genome and to precisely control the timing of gene expression, respectively. Such tools, in combination with FACS, transcriptome- and genome-wide sequencing, and other techniques will decipher how injury differentially affects the biology and behavior of certain cell populations in time and space. For instance, the mediation of the immediate local wound response could be studying by testing regeneration specific elements, such as putative enhancers identified by ATAC-seq, using simple fluorescent reporter lines. In parallel, the production of stem cell (neoblast) specific reporter lines should reveal the downstream effects of injury, ranging from cell differentiation, cell proliferation, to organogenesis. Altogether, our results show that *Hofstenia* will be a great system for insights into animal evolution and regeneration.

## Materials and Methods

### Animal culture

Adults *Hofstenia* were raised in plastic box containing 1 Liter of artificial filtered sea water (AFSW). Twice a week, boxes were cleaned, AFSW renewed, and animals were fed freshly-hatched artemia.

### Identification and cloning of regulatory elements of *Hofstenia* genes

*Hofstenia* full length transcript sequences of *troponin, sap* and *epiA* genes were identified in the *Hofstenia* transcriptome and the corresponding genomic loci were determined via BLAST (Gehrke *et al*., 2019). Primers were designed to amplify regions located 5’ to the ATG and 3’ to the STOP codon from genomic DNA. The plasmid from Renfer et al. (Renfer *et al*., 2010) was used as backbone for transgenesis. After digestion, *Hofstenia* promoters were directly cloned into the backbone. In our Kaede lines, 3’UTR regions were directly cloned into vectors already containing *Hofstenia* promoters and Kaede CDS. In our TagRFP-T lines, 3’UTRs were combined to the TagRFP-T CDS using the Gibson assembly Master Mix (Cat#E2611, New England BioLabs Inc). The resulting fragment was then amplified via PCR and digested prior ligation into a vector containing only *Hofstenia* promoter region. Following cloning, plasmid isolation and sequencing was performed to assess proper fragment insertion. For primer information, refer to the supplemental table 1.

### Injection solution

Prior to injection, 4 µg of the desired plasmid was fully digested with 4 µl of I-SceI enzyme, in a 50 µl reaction with 1X Cutsmart buffer, left 1 to 2 hrs at 37 °C, and followed by 20 minutes of heat inactivation at 65 °C. The digested plasmid was loaded in a 0.8% agarose (SeaKem® GTG™) gel. The fragment corresponding to the transgenic cassette was cut from the gel, purified with the NucleoSpin® Gel and PCR Clean-Up kit (Cat#740609, Mascherey-Nagel), eluted in 15-30 µl of elution buffer and quantified with a NanodropND-1000 spectrophotometer. The injection solution was prepared in a volume of 10 µl, by assembling the following reagents in nuclease-free water, in order to obtain the given final concentration: digested DNA (10-25 ng/µl); Fluorescein Dextran (D1820, Invitrogen; 1.25 µg/µl); 1X I-SceI Buffer (Renfer *et al*., 2010); I-SceI enzyme (Cat#R0694S, NEB; 0.375 U/µl). A quartz needle (Cat#QF100-70-10, Sutter Instrument) was back-loaded with 0.75 µl of the injection solution and manipulated with Narishige MMN1 and MO202U micro-manipulators. Injections were performed under a Leica MZ10F Stereomicroscope, using a Pico-liter Microinjector (Cat# PLI-90A, Warner Instruments).

### Embryo collection and microinjection dish

Embryos were collected with a glass pipette from culture boxes, then sorted under a stereomicroscope, in order to keep only the desired developmental stages (mainly zygotes and 2-cell stage embryos). Prior to each injection session, an injection dish was prepared using a plastic mold covered with square-base pins (side and length around 300 µm). The mold was deposited into a plastic culture dish (60 × 15 mm) half filled with melted agarose (1.2 % in MilliQ H2O) and left until full gelification, then gently removed, thus generating individual wells for the embryos. The injection dish was filled with AFSW and sorted embryos were transferred to the injection dish, then gently disposed into the wells for injection (one embryo per well). Following injection, embryos were collected from the wells with a glass pipette, transferred into a clean plastic dish, containing AFSW with Pennicillin and Streptavidin antibiotics (Cat#15140122, Thermofisher Scientific) at 5U/ml and placed in a 23 °C incubator. AFSW was only replaced 5 days after the injection.

### Transgenic embryo and animal care

Shortly prior hatching, embryos were screened for transgene expression by observing fluorescence with a Leica MZ10F stereomicroscope. Embryos exhibiting transgene expression were placed into individual wells within 24-well plates. After hatching, juveniles were transferred into new plates, with freshly made AFSW. From that point on until they reached adulthood, transgenic worms were transferred to clean plates and AFSW (without antibiotics), and fed (L-type *Brachionus* rotifers) twice a week. Over the course of their growth, animals were screened regularly to monitor transgene expression. Eventually, worms that were more likely to have integrated the transgene (based on brightness and area of expression) were placed into 6-well plates (each filled with 8 ml of AFSW), also with bi-weekly water renewals and feedings. For G0 (as well as for the following generations) animals, sexually mature worms were crossed with wild type animals or with other transgenic worms. Their progeny were screened and taken care of as described above.

### Immunohistochemistry (IHC), FISH and imaging

Animals were relaxed for five minutes in AFSW containing 0.5 mg/ml of Tricaine and with adjusted pH (8-8.5). This solution was then replaced by the fixation solution: 4% Paraformaldehyde (PFA) in AFSW, with 0.05% Triton X-100. Animals were fixed either 3hrs at room temperature (RT) or overnight (O/N) at 4°C, with constant agitation. Following fixation, FISH procedure was performed as described previously (Srivastava *et al*., 2014). For IHC, fixed worms were briefly washed three times in 1X Phosphate Buffer Saline with 0.1% Triton X-100 (PBSTx), then blocked for 2 hrs at RT in PBSTx with 1% Bovine Serum Albumin (BSA). Primary antibody incubation was performed in renewed blocking solution, at 4°C, during 48-72 hrs, with constant agitation. Long washes in PBSTx were repeated 5-6 times along the day, before incubation (48-72 hrs) with secondary antibodies, in PBSTx. Samples were then washed extensively in PBSTx, and counterstained with 4′,6-diamidino-2-phenylindole (DAPI), (268298, EMD Millipore), at 2 µg/ml, for 2 hrs, at RT. Primary antibodies used: anti-TagRFP (Cat#R10367, Invitrogen) at the dilution of 1:1000, custom made anti-*Hm*Tropomyosin at the concentration of 1 µg/ml. Secondary antibodies used : Goat anti-rabbit, Alexa Fluor® 568 (Cat#ab175471, Abcam); Goat anti-rabbit, Alexa Fluor® 488 (Cat#ab150077, Abcam). Imaging was performed with a Leica SP8 confocal microscope. Post-acquisition treatment of confocal images was performed with Fiji; in particular, 3D rendering images were made with the 3D Viewer plugin.

### Regeneration assay

One month old G2 *epiA*::*TagRFP-T* transgenic worms were relaxed in AFSW with 0.5 mg/ml Tricaine, then bisected with a razor blade at the level of the middle stripe. Following amputation, fragments were rinsed 3 times in AFSW without Tricaine, then placed into AFSW within 12-well plates, and left to regenerate at 23°C in an incubator. AFSW was changed daily until fixation in 4% PFA for imaging.

### FACS assay and Immunocytochemistry (ICC)

Cell dissociation of transgenic and wild type worms was done by placing one adult worm in a 2ml tube, replacing AFSW with calcium- and magnesium-free AFSW (CMFSW) with 1% horse serum, and pipetting until full dissociation. Dissociated cells were filtered using a 40 µm cell strainer into a clean 2ml tube, then cleaned from debris by centrifugation (5 min., 500 g, at 4 °C) over a 4% BSA cushion in CMFSW. The resulting pellet was resuspended into CMFSW with 1% horse serum and placed on ice before labeling with Hoescht 33342 (Cat#H3570, Invitrogen; 10 mg/ml final) and propidium iodide (Cat# P3566, Invitrogen; 1 mg/ml final) for 30 min. Sorting was done using a Beckman Coulter MoFlo Astrios EQ Cell Sorter. For ICC, sorted cells were collected into 2ml tubes with 1 ml of CMFSW and centrifuged (5 min., 500 g, at 4 °C). The pellets were resuspended in 150 µl of CMFSW, then deposited onto lysine-coated coverslips, at the bottom of a 12-well plate (from that point on, all incubations were performed in the dark), and let to adhere for 1 hour. Fixation was performed for 20 min. with 4% PFA in CMFSW and followed by 3 quick washes with PBSTx. Samples were blocked with PBSTx + 10% horse serum, then incubated with primary antibody (2 µg/ml) in block, 2 hours at RT. After 3 PBSTx washes, samples were incubated with the secondary antibody for 2 hours, at RT. Finally, samples were washed 3 times with PBSTx, incubated in Hoescht 33342 (1 mg/ml) and mounted on microscopy slides for imaging.

### Live imaging of embryos

Embryos were placed into Nunc™ bottom glass dishes (Cat#150682, Thermo Scientific™) containing 4 ml of AFSW and imaged with a Leica SP8 confocal. For time-lapse imaging, Pennicillin and Streptavidin antibiotics (1 Cat#5140122, Thermofisher Scientific) at 5U/ml were added to the water.

### CRISPR/CAS9 genome editing

A target site for CAS9 enzyme was identified with Geneious (https://www.geneious.com) in the *Hofstenia* GRP78 locus. Primers were designed in order to synthesize a single guide RNA (sgRNA) corresponding to this target site, using the MEGAscript™ T7 Transcription Kit (Cat#AM1334, Invitrogen™) (Zhang and Reed, 2016). sgRNA was then purified with phenol-chloroform followed by isopropanol / ammonium acetate precipitation, and resuspended in nuclease free water. Embryos were injected with a mixture of sgRNA at 600 ng/µl, CAS9 enzyme at 1µg/µl (Cat#1074181, Integrated DNA Technologies) and fluorescein dextran, then left to develop until pre-hatchling stage. Genomic DNA from individual embryos was isolated with the NucleoSpin® Tissue XS kit (Cat#740901, Macherey-Nagel). A PCR reaction was performed using the isolated genomic DNA with primers flanking the CAS9 target site. The T7E1 assay (Cat#M0302, NEB) was then applied as in (Sato *et al*., 2018) to the PCR product (half of the product being treated with T7 endonuclease, in the other half, T7E1 was replaced by water) which was run on an agarose gel to assess for multiple band detection. Control bands were then extracted from the gel and sequenced in order to assess for the presence of indels at the target site.

### Resource Availability

#### Lead Contact

Further information and requests for resources and reagents should be directed to and will be fulfilled by the Lead Contact, Mansi Srivastava.

#### Materials Availability

Plasmids generated in this study will be deposited to Addgene. Transgenic lines generated in this study will be available from the Lead Contact with a completed Materials Transfer Agreement.

## Acknowledgements

We thank Dr. Mark Martindale for guidance on embryonic microinjections and Dr. Seth Donoughe for help with making injection molds. L.R. and M.S. are supported by the Searle Scholars Program, the Smith Family Foundation, and NIH (1R35GM128817)

## Author Contributions

L.R. and M.S. designed the study and wrote the manuscript. L.R. conducted the experiments.

## Supplemental Information

**Supp. Fig. 1:**
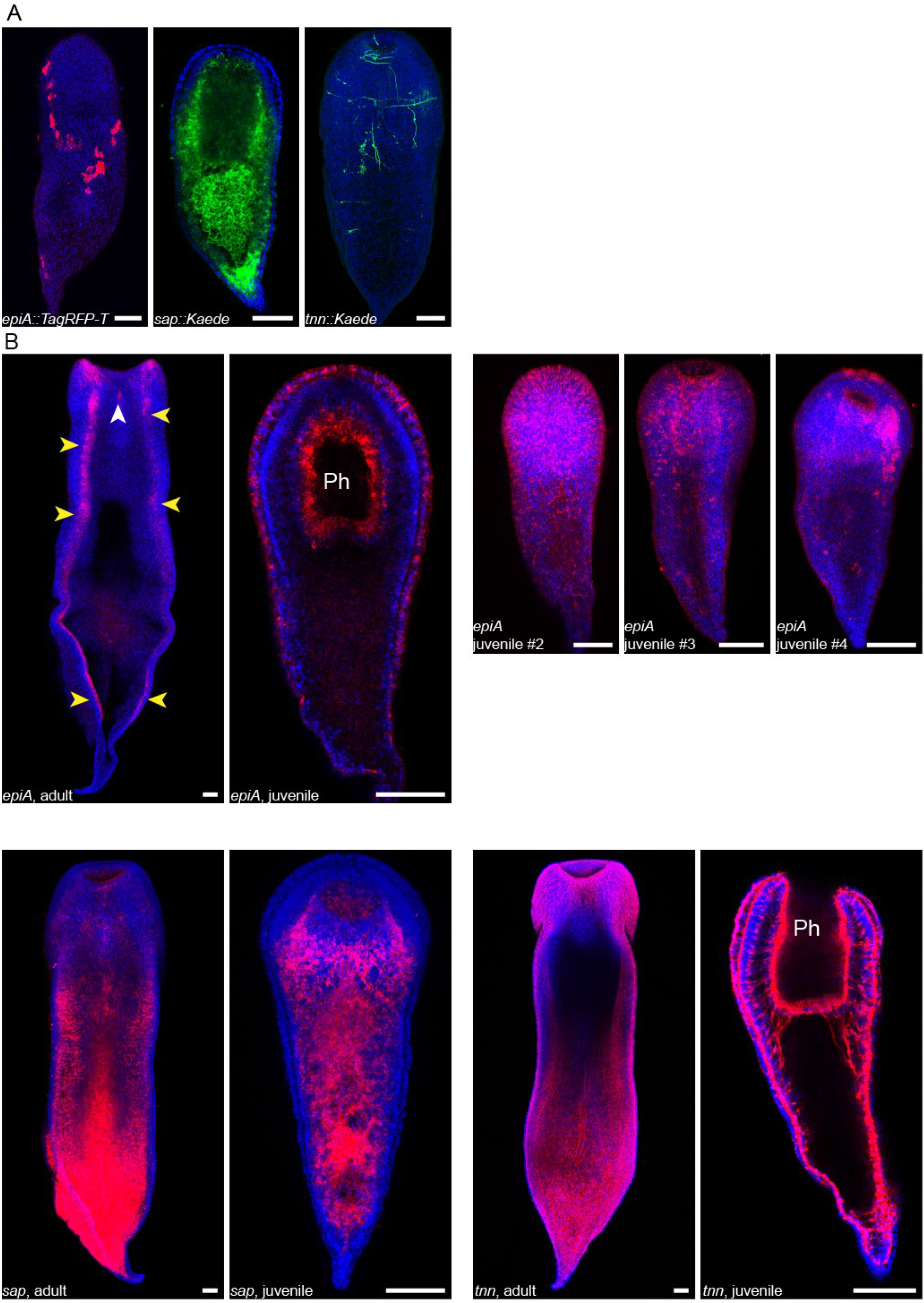
Expression patterns for wild-type and transgenic *epi-a, sap* and *tnn* genes. **(A)** Mosaic transgene expression in juvenile worms. Transgene name is indicated in the bottom left corner. TagRFP-T (red); Kaede (green); DAPI (nuclei, blue). **(B)** Fluorescent *in situ* hybridization of riboprobes (red) for *epiA, sap* and *tnn* are shown in adults and hatchlings of *Hofstenia* worms. White arrowhead points to *epiA*-expressing cells in the male penis stylet opening. Yellow arrowheads point at *epiA*-expressing cells in adult that were not consistently recapitulated by transgene expression. An example of the variety of expression patterns observed for *epiA* shown in four different hatchlings. All worms are oriented anterior side up and images of adult worms show their ventral side. Nuclei are counterstained with DAPI (blue). *Ph, pharynx lumen*. Scale bars, 100 µm.

**Supp. Fig. 2:**
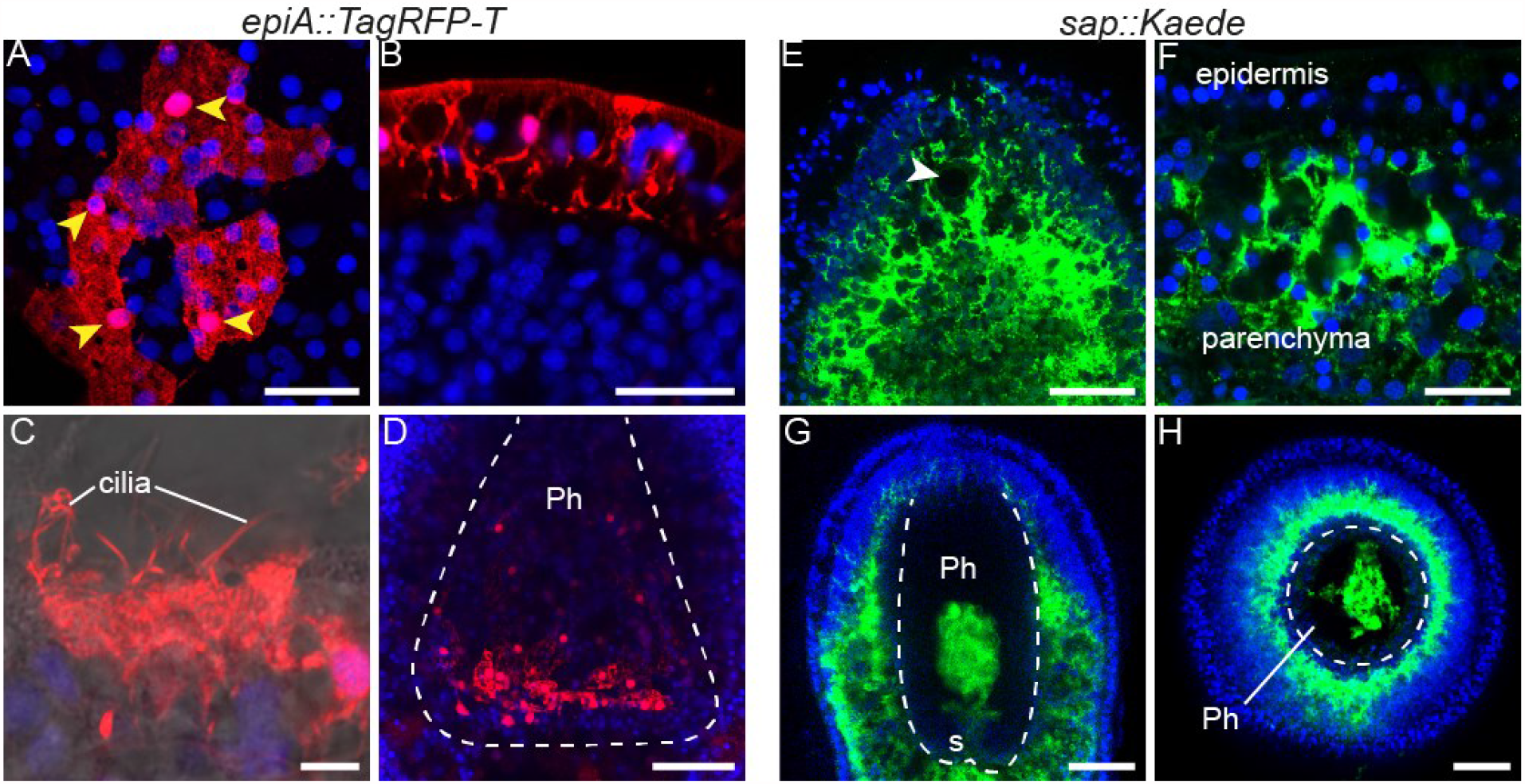
Transgene expression reveals details of acoel anatomy. Expression of *epiA::TagRFP-T* (A-D) and *sap::Kaede* (E-H) transgenes in hatchlings. Top (**A**) and side view **(B)** of a patch of epidermal cells. Yellow arroheads point at epidermal cell nuclei positions. (**C**) Ciliature of an epidermal cell. (D) *EpiA*+ cells, lie on the pharynx wall. (**E**) Kaede expressing cells in a hatchling head (white arrowhead indicates the statocyst position). (**F**) Kaede expressing cells at the parenchyma-epidermis junction. (**G, H**) Kaede+ digestive tissue protruding, through the pharynx sphincter, within the pharynx cavity, as seen in longitudinal (G) and cross (H) sections. White dashed lines delineate the pharynx cavity; *Ph, pharynx lumen; s, pharynx sphincter*. TagRFP-T (red); Kaede (green); DAPI (nuclei, blue). Scale bars, A, B, F, 20 µm; C, 5 µm; D, E, G, H, 50 µm.

**Supp. Fig. 3:**
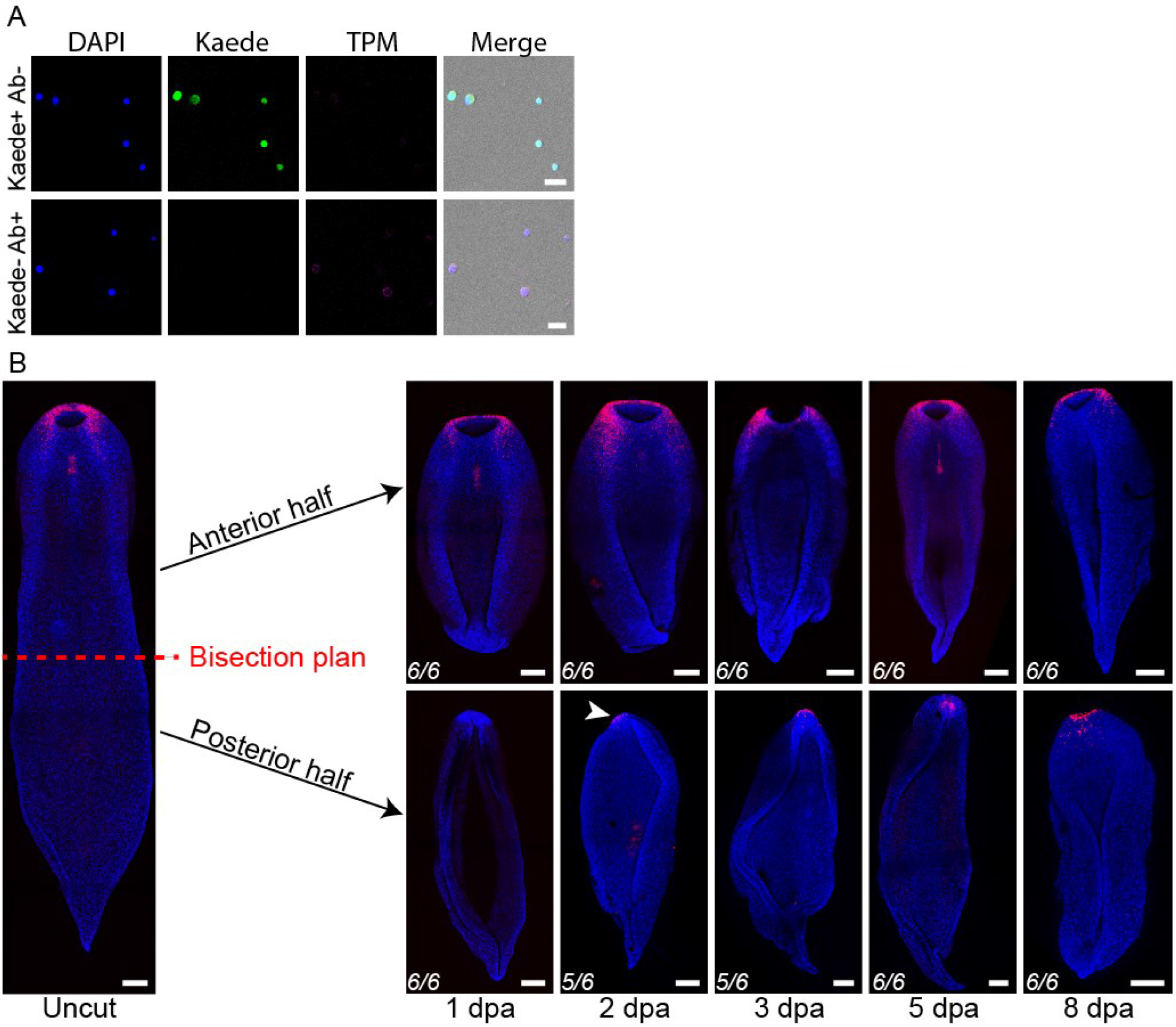
Transgenic toolkit. **(A)** Controls for FACS and ICC. Anti-Tropomyosin (TPM) immunohistochemistry on FACS-isolated cells, from *tnn::Kaede* transgenics. Scale bars, 20 µm. (**B**) Regeneration in *epiA::TagRFP-T* transgenics. Regeneration time course of both anterior and posterior fragments after sub-pharyngeal bisection. Time elapsed since amputation is indicated at the bottom (dpa, days post amputation). White arrowhead points to *de novo* transgene expression in posterior fragment. For each time points, 6 worms were amputated, the frequency of the phenotype is indicated in the left bottom corner. All images are projection of confocal z-stacks, taken ventrally, and oriented anterior side up. Scale bars, 200 µm.

**Supp Fig. 4:**
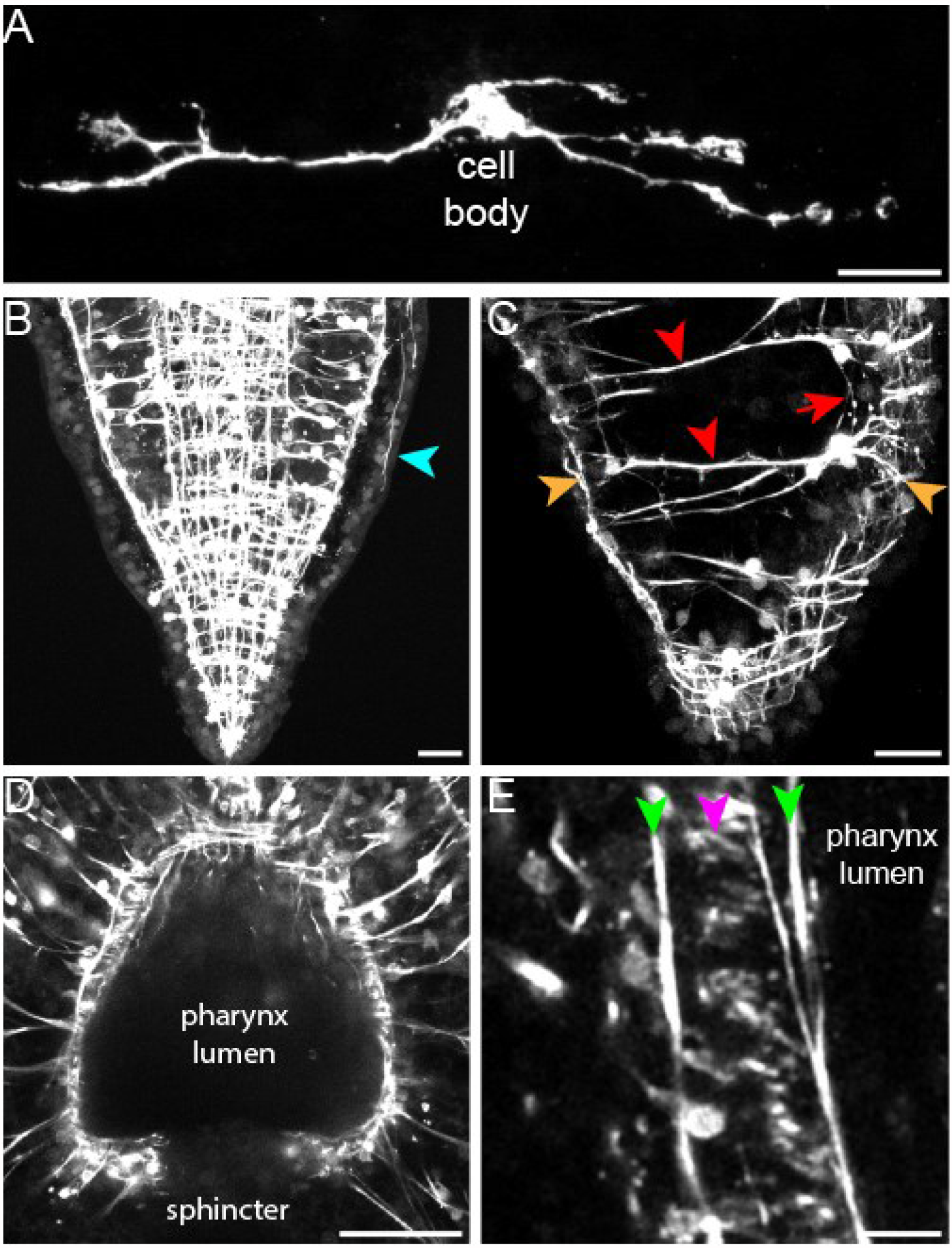
Details of muscle anatomy. (**A**) Full length peripheral muscle fiber in G0 animal. **(B)** The tip of the tail appears deprived of peripheral muscles. Blue arrowhead points to the posterior extremity of a peripheral muscle. (**C**) Posterior parenchymal muscles (red arrowhead) connect body wall fibers together (orange arrowheads). Cellular projections can be observed connecting parenchymal fibers together (red arrow). (D, E) The pharynx is contained within a dense, basket-shaped (**D**), three-layered (**E**) muscular network. Green arrowheads indicate longitudinal pharyngeal muscles; magenta arrowhead point at circular pharyngeal muscles. Scale bars, A, E,10 µm; B, C, 20 µm; D, 50 µm.

**Supp. Fig. 5:**
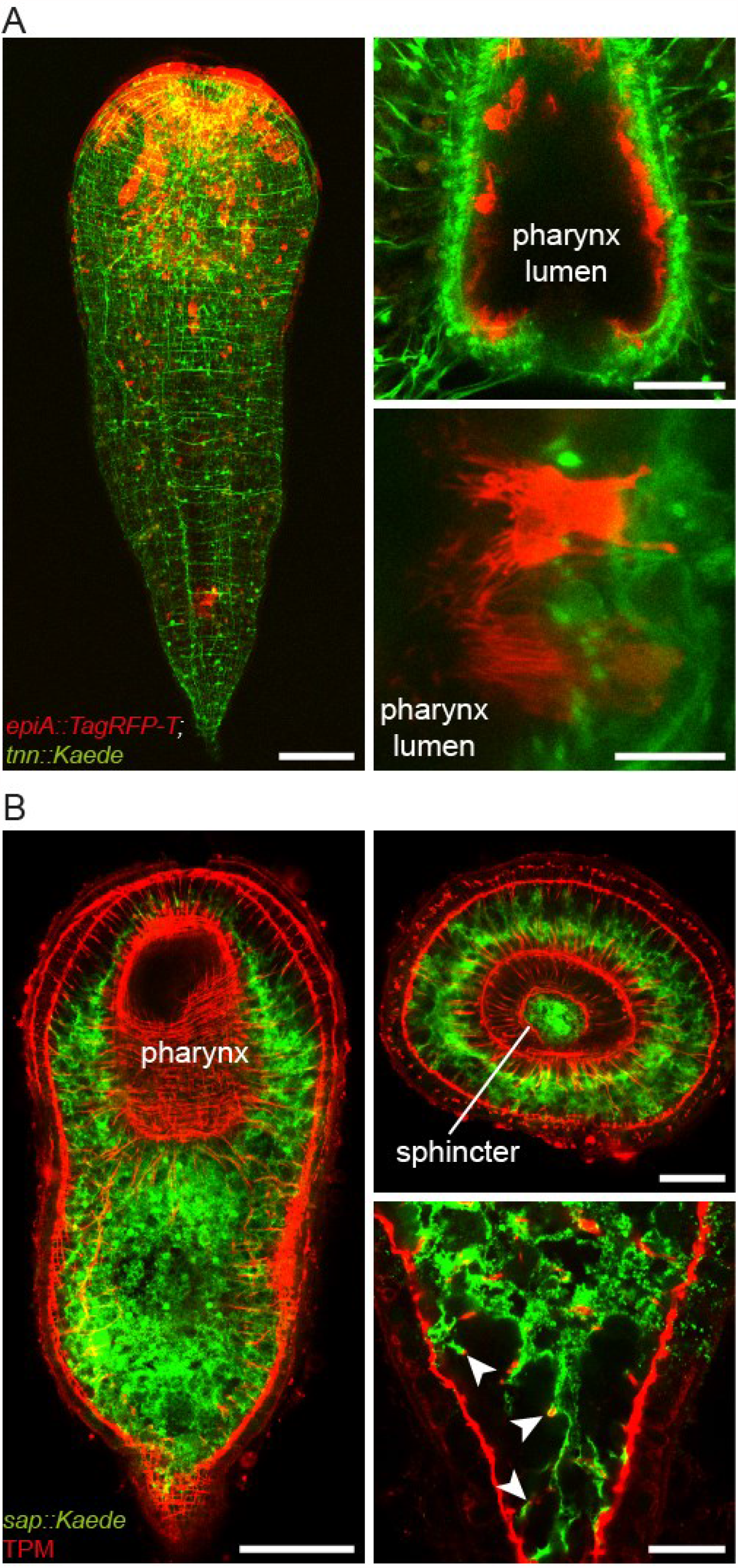
Dual labeling of muscle with *epiA*+ and *sap*+ cells. (**A**) Crossing of G1 *epiA::TagRFP-T* with G1 *tnn::Kaede* transgenic lines. (**Left**) Double transgenic *epiA::Tag-RFP-T* ; *tnn::Kaede*, hatchling, projection of confocal z-stack. (**Right, top**) Pharynx, longitudinal section. (**Right, bottom**) Ciliature of *epiA*+ pharingeal cells. Kaede (green); TagRFP-T (red). Scale bars, A, 100 µm; B, 50 µm; C, 10 µm. (**B**) Dual abeling of muscle with *sap*+ cells. Anti-TPM immunohistochemistry on *sap::Kaede* G2 transgenic animals. (**Left**) Whole worm (hatchling), projection of confocal z-stack. (**Right, top**) Pharynx, posterior, cross section. (**Right, bottom**) Association of digestive tissue with muscle fibers (white arrowheads), in the tail of the animal. Kaede (green); TPM (red). Scale bars, A, 100 µm; B, 50 µm; C 20 µm.

**Supp. Fig. 6:**
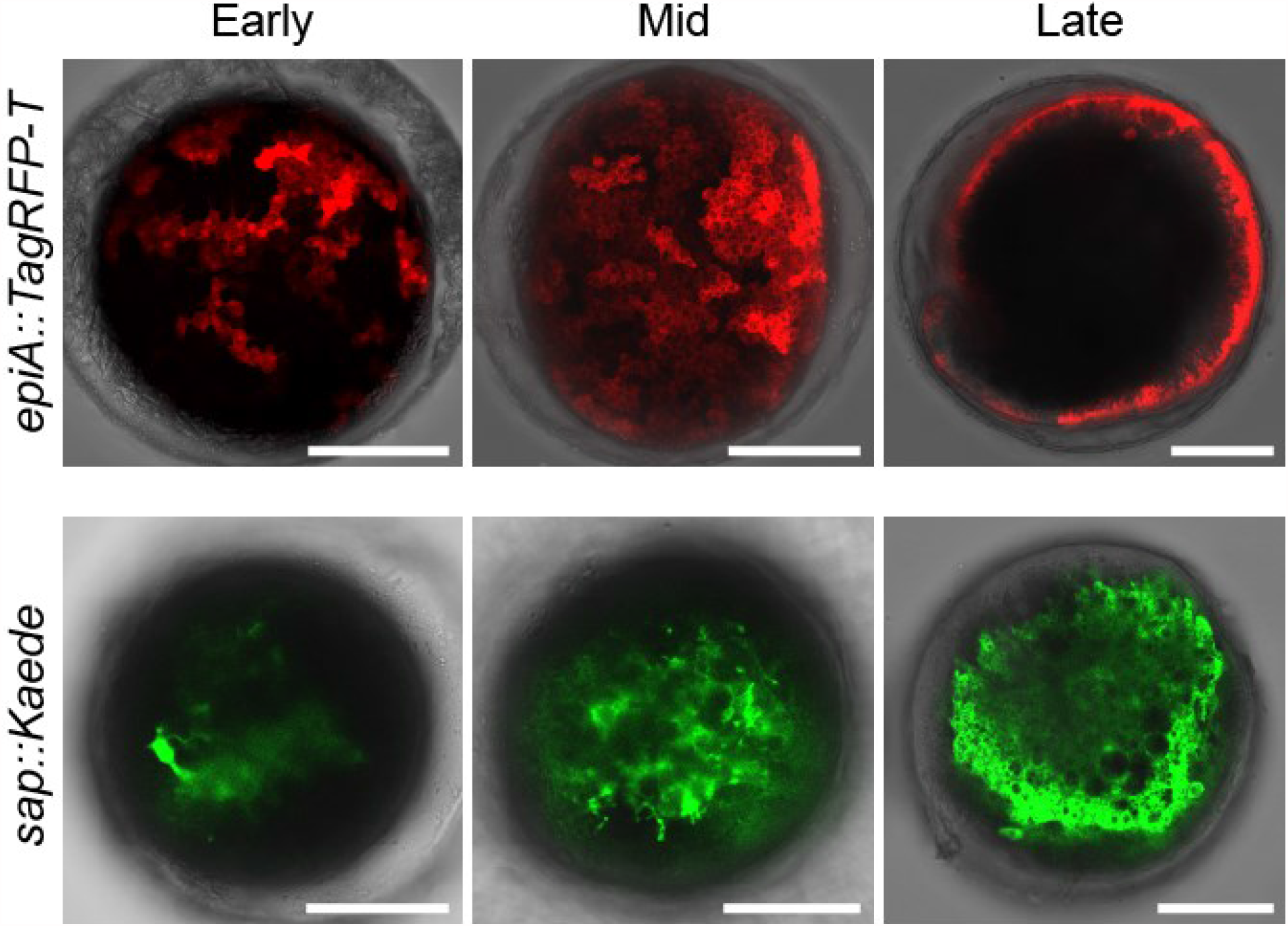
Transgenesis enables live studies of organogenesis. Transgene expression as seen in G1 or G2 embryos at three different time points : right after detection (Early, 4.5 days post fertilization), then 1-2 days later (Mid) and finally, prior hatchling (Late), when the embryos are constantly spinning within their capsule. Name of the transgene is indicated on the left side of the panel. Scale bars, 100 µm.

**Supp. Fig. 7:**
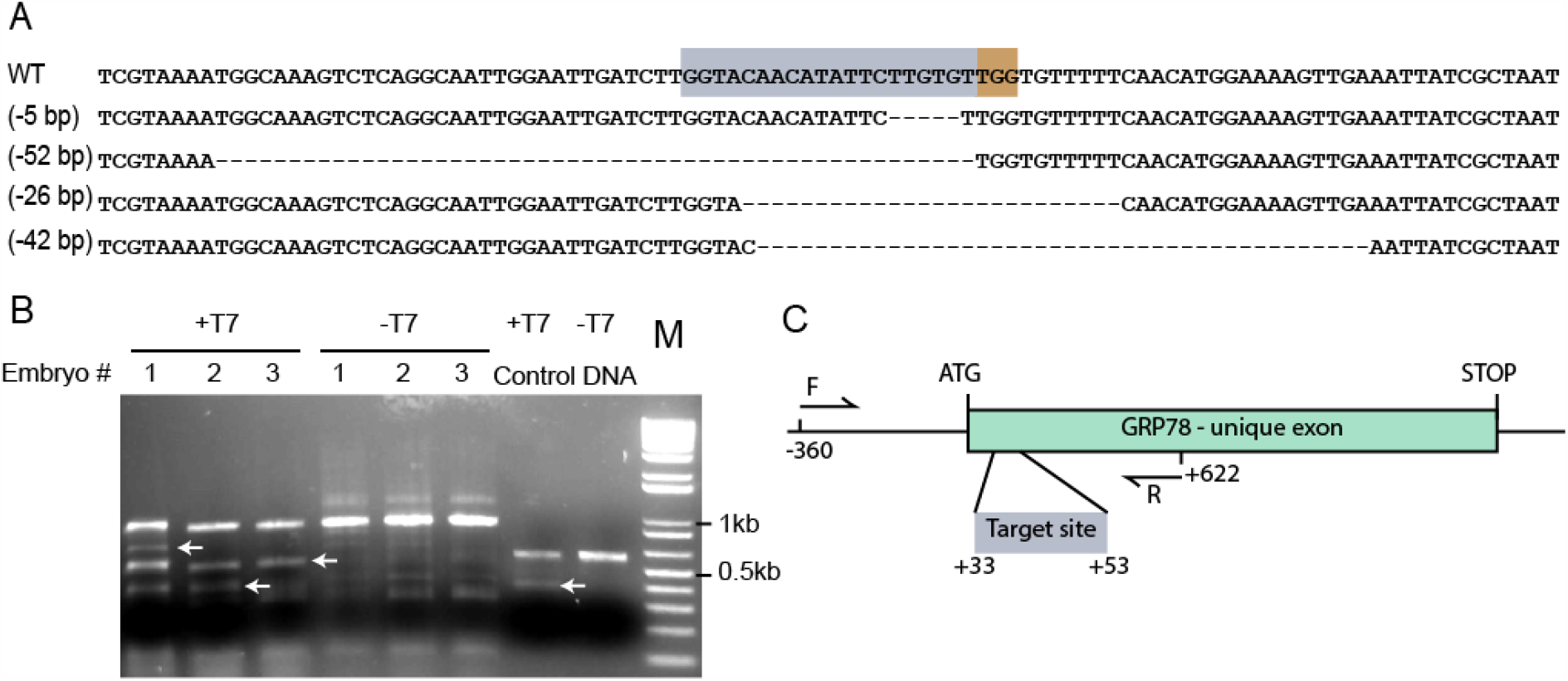
Genome editing. (**A**) Sequences of the CAS9 targeted GRP78 locus, from 4 different injected embryos. Wild-type DNA sequence for the locus is shown on top; the number of deleted bases is indicated between parenthesis; grey and brown boxes show CAS9 target and PAM sites, respectively. (**B**) Gel migration of PCR amplified GRP78 locus with (+T7) or without (-T7) T7E1 treatment. White arrows show bands smaller than the expected amplicon, due to T7 activity on double strand DNA mismatch resulting from CRISPR/CAS9 indels (6 out of 9 injected embryos showed similar gel bands and indels); M, markers of molecular weight. **(C)** GRP78 locus, with the CAS9 target site and the primers used for pcr amplification.

**Supp. Video 1 : Morphology of epidermal cells**. 360° rotation of a 3D image, reconstructed from confocal z-stack, showing a patch of *epiA::TagRFP-T* expressing cells in a G0 fixed juvenile. White arrowhead points to cilia, on the apical side; white arrow points to the epidermal processes.

**Supp. Videos 2A and 2B : Interlocking of parenchymal and body wall muscle fibers**. 360° rotation, along the X (**2A**) and Y (**2B**) axis, of a 3D image, reconstructed from confocal z-stack of a G2 *tnn::Kaede* juvenile worm. Colored arrows point to the different muscle categories. *PA, PE, BW, parenchymal, peripheral and body wall muscles, respectively*.

**Supp. Videos 3A and 3B : Interlocking of single parenchymal and body wall muscle fibers**. 360° rotation, along the X (**3A**) and Y (**3B**) axis, of a 3D image, reconstructed from confocal z-stack of a G0 *tnn::TagRFP-T* juvenile worm. *PA, BW, parenchymal,and body wall muscles, respectively*.

**Supp. Video 4 : Emergence of the musculature during embryogenesis**. Time-lapse imaging of a G2 *tnn::Kaede* embryo over 63hrs of development, from 3.5 to 6 days post laying. Time stamp, *hh::mm::ss*.

**Supp. Video 5 : Interaction of posterior epidermis with body wall muscle**. 360° rotation of a 3D image, reconstructed from confocal z-stack, showing a patch of *epiA::TagRFP-T* (red) expressing cells over the body wall muscles, revealed by anti-TPM IHC (green) in a G1 *epiA::TagRFP-T* juvenile worm. Nuclei are counterstained with DAPI (blue).

